# Comparative study of transcriptomics-based scoring metrics for the epithelial-hybrid-mesenchymal spectrum

**DOI:** 10.1101/2020.01.02.892604

**Authors:** Priyanka Chakraborty, Jason T George, Shubham Tripathi, Herbert Levine, Mohit Kumar Jolly

**Author notes:** Corresponding author (M.K.J.).

## Abstract

The Epithelial-mesenchymal transition (EMT) is a cellular process implicated in embryonic development, wound healing, and pathological conditions such as cancer metastasis and fibrosis. Cancer cells undergoing EMT exhibit enhanced aggressive behavior characterized by drug resistance, tumor-initiation potential, and the ability to evade immune system. Recent *in silico, in vitro*, and *in vivo* evidence indicates that EMT is not an all-or-none process; instead, cells stably acquire one or more hybrid epithelial/mesenchymal (E/M) phenotypes which often can be more aggressive than purely epithelial or mesenchymal cell populations. Thus, the EMT status of cancer cells can prove to be a critical estimate of patient prognosis. Recent attempts have employed different transcriptomics signatures to quantify EMT status in cell lines and patient tumors. However, a comprehensive comparison of these methods, including their accuracy in identifying cells in the hybrid E/M phenotype(s), is lacking. Here, we compare three distinct metrics that score EMT on a continuum, based on the transcriptomics signature of individual samples. Our results demonstrate that these methods exhibit good concordance among themselves in quantifying the extent of EMT in a given sample. Moreover, scoring EMT using any of the three methods discerned that cells undergo varying extents of EMT across tumor types. Separately, our analysis also identified tumor types with maximum variability in terms of EMT and associated an enrichment of hybrid E/M signatures in these samples. Moreover, we also found that the multinomial logistic regression (MLR) based metric was capable of distinguishing between ‘pure’ individual hybrid E/M vs. mixtures of epithelial (E) and mesenchymal (M) cells. Our results, thus, suggest that while any of the three methods can indicate a generic trend in the EMT status of a given cell, the MLR method has two additional advantages: a) it uses a small number of predictors to calculate the EMT score, and b) it can predict from the transcriptomic signature of a population whether it is comprised of ‘pure’ hybrid E/M cells at the single-cell level or is instead an ensemble of E and M cell subpopulations.

## Introduction

The Epithelial-mesenchymal transition (EMT) is a cell biological process crucial for various aspects of tumor aggressiveness - cancer metastasis (Jolly et al., 2017), resistance against cell death (Huang et al., 2013), metabolic reprogramming (Thomson et al., 2019), refractory response to chemotherapy and radiotherapy (Kurrey et al., 2009), tumor-initiation potential (Jolly et al., 2014), and immune evasion (Tripathi et al., 2016; Terry et al., 2017) - thus eventually affecting patient survival (Tan et al., 2014). EMT is a multidimensional, nonlinear process that involves changes in a compendium of molecular and morphological traits, such as altered cell polarity, partial or complete loss of cell-cell adhesion, and increased migration and invasion. Cells may take different routes in this multidimensional landscape as effectively captured by recent high-throughput dynamic approaches (Karacosta et al., 2019; Watanabe et al., 2019). Initially thought of as binary, EMT is now considered as a complex process involving one or more hybrid epithelial/mesenchymal (E/M) states (Jolly and Celia-Terrassa, 2019). These hybrid E/M states can be more plastic and tumorigenic than ‘purely epithelial’ or ‘purely mesenchymal’ ones, thus constituting the ‘fittest’ phenotype for metastasis (Grosse-Wilde et al., 2015; Bierie et al., 2017; Pastushenko et al., 2018; Kröger et al., 2019; Tripathi et al., 2019b). Consequently, the presence and frequency of such hybrid E/M cells in primary tumors and in circulating tumor cells (CTCs) can be associated with poor patient survival (Jolly et al., 2019a; Saxena et al., 2019). Computational methods aimed at quantifying EMT on a continuous spectrum in order to enhance diagnostic, prognostic, and therapeutic intervention are therefore indispensable.

Various methods have been developed to obtain a quantitative measure of the extent of EMT (hereafter, referred to as EMT score) that cells in a given sample have undergone. Here we focus on methods accomplishing this task using the gene expression data. First, a 76-gene EMT signature (76GS; hereafter referred to as 76GS method) was developed and validated using gene expression from non-small cell lung cancer (NSCLC) cell lines and patients treated in the BATTLE trial (Byers et al., 2013). This scoring method calculates EMT scores based on a weighted sum of the expression levels of 76 genes; the weight factor of a gene is the correlation coefficient between the expression level of that gene and that of CDH1 (E-cadherin) in that dataset; thus, the absolute EMT scores of epithelial samples using the 76 GS method are relatively higher than those of mesenchymal samples (Guo et al., 2019). Second, an EMT score separately for cell lines and tumors was developed based on a two-sample Kolmogorov-Smirnov test (KS; hereafter referred to as the KS method). This score varies on a scale of −1 to 1, with the higher scores corresponding to more mesenchymal samples (Tan et al., 2014). Third, a multinomial logistic regression (MLR; hereafter referred to as the MLR method) based model quantified the extent of EMT in a given sample on a scale of 0 to 2. This method particularly focuses on characterizing a hybrid E/M phenotype using the expression levels of 23 genes – 3 predictors and 20 normalizers – identified through NCI-60 gene expression data. It consequently calculates the probability that given sample belongs to epithelial, mesenchymal, or hybrid E/M categories. An EMT score is assigned based on those probabilities; the higher the score, the more mesenchymal the sample is (George et al., 2017). A comparative analysis of these methods in terms of similarities, differences, strengths, and limitations, remains to be done.

Here, we present a comprehensive evaluation of these methods – 76GS, KS and MLR - in terms of quantifying EMT and characterizing the hybrid E/M phenotype. First, we calculate the correlations observed across different *in vitro, in vivo*, and patient datasets, and observe good quantitative agreement among the scores calculated using these 3 methods. This analysis suggests that all of them, despite using varied gene lists and methods, concur in capturing a generic trend, embedded in the multi-dimensional EMT gene expression landscape. Second, we identify which cancer types are more heterogeneous than others in terms of their EMT status; intriguingly, our results show that enrichment for a hybrid E/M phenotype contributes to heterogeneity. Third, we compare the ability of these methods to distinguish between ‘pure’ individual hybrid E/M cells vs. mixtures of epithelial (E) and mesenchymal (M) cells that can exhibit an EMT score similar to that of hybrid E / M samples. Our results offer proof-of-principle that the MLR method can identify these differences. Overall, our results demonstrate the consistency of these EMT scoring metrics in quantifying the spectrum of EMT. Moreover, two advantages of MLR method are highlighted – namely, the use of a small number of predictors to calculate the EMT score, and the ability to characterize difference between admixtures of E and M cells vs. truly hybrid E/M cells.

## Methods and Materials

### Software and datasets

All computational and statistical analyses were performed using R (version 3.4.0) and Bioconductor (version 3.6). Microarray datasets were downloaded using *GEOquery* R Bioconductor package (Davis and Meltzer, 2007). TCGA datasets were obtained from the *UCSC xena tools* (Wang et al., 2019a). C CLE and NCI60 datasets were downloaded from respective websites.

### Preprocessing of microarray data sets

All the microarray datasets were preprocessed to obtain the gene-wise expression for each sample from probe-wise expression matrix. To map the probes to genes, relevant platform annotation files were utilized. If there were multiple probes mapping to one gene, then the mean expression of all the mapped probes were considered for that gene.

### Calculation of EMT scores

EMT scores were calculated for samples in a particular data set using all three methods. For a particular microarray data set, expression of respective gene signatures was given as an input to calculate EMT score using all three different methods.

#### 76GS

The EMT scores were calculated based on a 76-gene expression signature reported (Byers et al., 2013) (**Table S1**) and the metric mentioned based on that gene signature (Guo et al., 2019). For each sample, the score was calculated as a weighted sum of 76 gene expression levels and the scores were centered by subtracting the mean across all tumor samples so that the grand mean of the score was zero. Negative scores can be interpreted as mesenchymal phenotype whereas the positive scores as epithelial.

#### MLR

The ordinal MLR method predicts EMT status based on the order structure of categories and the principle that the hybrid E/M state falls in a region intermediary to E and M. Quantitative estimates of EMT spectrum were inferred based on the assumptions and equations mentioned (George et al., 2017) (**Table S2**). The samples are scored ranging from 0 (pure E) to 2 (pure M), with a score of 1 indicating a maximally hybrid phenotype. These scores are calculated based on the probability of a given sample being assigned to the E, E/M and M phenotype.

#### KS

The KS EMT scores were calculated as previously reported (Tan et al., 2014) (**Table S3, S4**). This method compares cumulative distribution functions (CDFs) of epithelial and mesenchymal signatures. First, the distance between epithelial and mesenchymal signatures was calculated via the maximum distance between their CDFs as follows: For CDFs *F*_*E*_ (*x*) and *F*_*M*_ (*x*) representing the levels of transcript *x* for epithelial and mesenchymal signatures, respectively, the distance between signatures is assessed by using the uniform norm

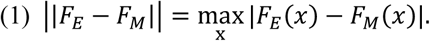

This quantity represents the test statistic in the subsequent two-sample test used to calculate the EMT score. The score is determined by hypothesis testing of two alternative hypotheses as follows (with the null hypothesis being that there is no difference in CDF of mesenchymal and epithelial signatures): (1) CDF of mesenchymal signature is greater than CDF of epithelial signature. (2) CDF of epithelial is greater than CDF of mesenchymal signature. Sample with a positive EMT score is mesenchymal whereas negative EMT score is associated with epithelial phenotype.

### Correlation analysis

Correlation between EMT scores was calculated by Pearson’s correlation, unless otherwise mentioned.

### Survival analysis

All samples were segregated into EMT^high^ and EMT^low^ groups based on the mean values of EMT score. Observed survival distributions are graphically depicted for the abovementioned two categories using Kaplan–Meier plots.

### Mixture curve analysis

For each dataset analyzed using mixture curves, the most mesenchymal (pure-M) and most epithelial (pure-E) samples were identified by ordering samples based on MLR EMT score and selecting the top and bottom 35 samples, respectively. The mean or median was calculated for the pure-E and pure-M samples as a representative of the purified E or M state in the MLR predictor space. From this, the mixture curve is derived by taking all convex combinations of purified states. Individual samples within a given dataset were ranked based on their proximity to the mixture curve using the usual l2-norm distance. The top 10, 20, 50, and 100 samples closest to, and furthest from, the mixture curve were used as representative mixtures of E and M populations and hybrid E/M signatures, respectively.

## Results

### Concordance in capturing EMT response

We used three different EMT scoring methods to quantify the extent of EMT in given transcriptomics data; each method utilizes a distinct gene set as well as a different underlying algorithm. In the 76GS method, the higher the score, the more epithelial a sample is, given that the method calculates as weighted sum of expression levels of 76 genes, with the weight factor being correlation coefficient with levels of canonical epithelial marker CDH1 (**Fig 1A**). This method has no specific pre-defined range of values, although the range of values obtained are bounded by the maximal possible value of gene expression detected by microarray. Unlike the 76 GS method, the MLR and KS methods have predefined scales for EMT scores. MLR and KS score EMT on a spectrum of [0, 2] and [-1, 1] respectively, with higher scores indicating mesenchymal signatures (**Fig 1B, C**). While MLR and KS methods are absolute, requiring a fixed transcript signature for EMT score calculation, the 76GS method of EMT scoring depends on the number and nature of samples analyzed in a given dataset. Consequently, a hybrid E/M sample may have a (pseudo) high 76GS score whenever the available dataset contains more mesenchymal samples, or a (pseudo) low score for datasets enriched in epithelial samples. Each scoring method also varies in the number of required gene transcripts: while the MLR method utilizes 23 entries, the 76GS methods requires 76. The KS method utilizes 315 and 218 transcripts for tumor samples and cell lines samples, respectively.

**Figure 1:**
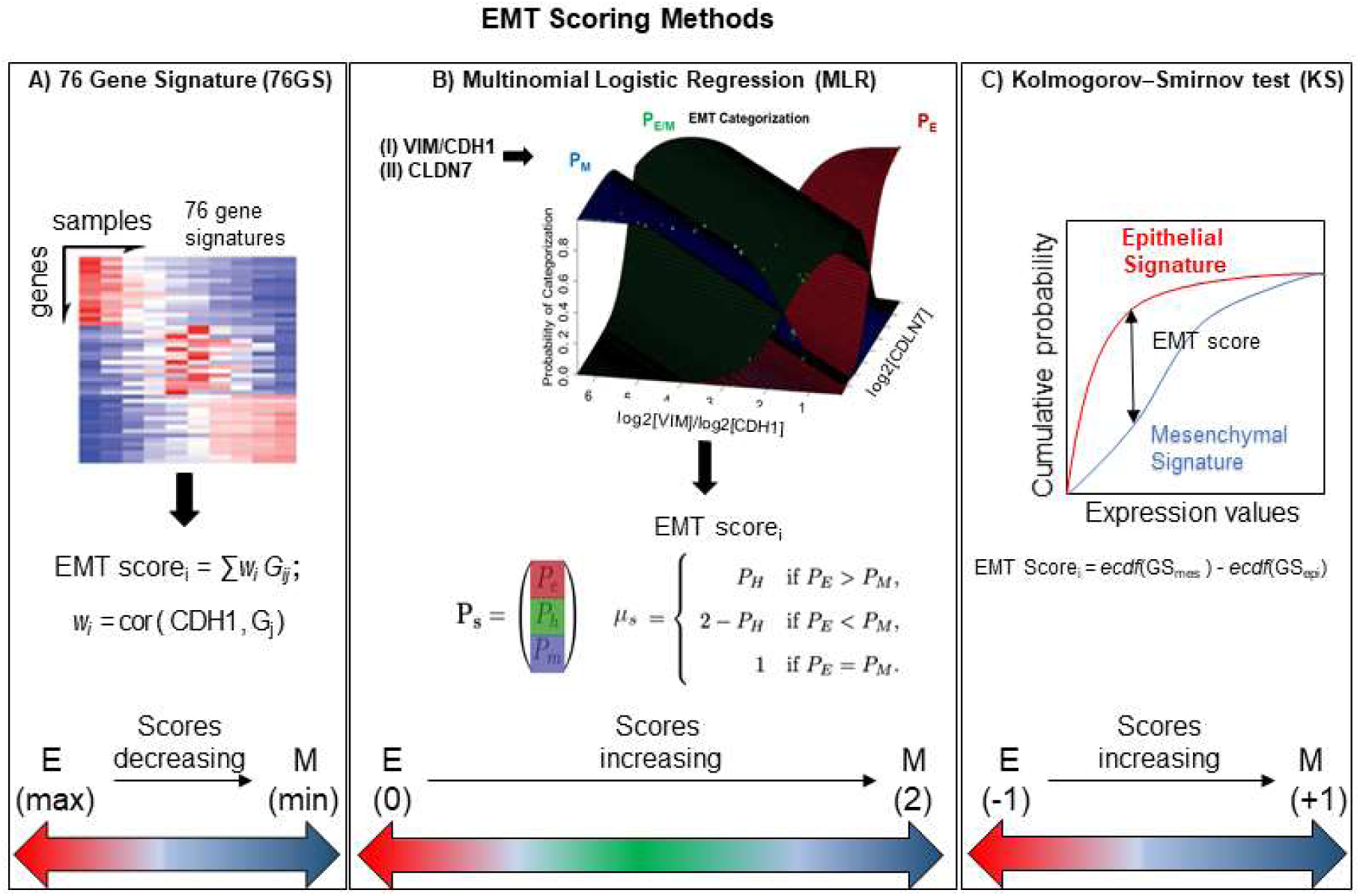
General outline of all three EMT scoring methods. **A)** 76GS score is calculated by weighted sum of 76 genes, where EMT score_i_ is score for i^th^ sample, w_j_ is correlation of j^th^ gene (G_j_) with CDH1 gene in that dataset to which the sample i belongs, G_ij_ is the j^th^ gene’s normalized expression in i^th^ sample. **B)** MLR utilizes log2(VIM)/log2(CDH1) and log2(CLDN7) space to predict categorization of a sample into E, E/M or M category. Where, P_E_, P_H_, and P_M_, are the probabilities of a sample falling into each phenotype. EMT score_i_ is the score for i^th^ sample, which is defined in relation to P_E_, P_H_, and P_M_.**C)** KS score is estimated by the empirical cumulative distributions of Epithelial and Mesenchymal gene set, denoted by ecdf (GS_mes_) and ecdf (GS_epi_) respectively. EMT score _i_ is the maximum vertical distance between the ecdf (GS_mes_) and ecdf (GS_epi_), (given by Eq. 1 in Methods section) for a given sample i.

We first investigated the extent of concordance in EMT scores calculated via these three methods for well-studied cohort of cancer cell lines: NCI-60 and CCLE (Shankavaram et al., 2009; Barretina et al., 2012). We expected to see a negative correlation between EMT scores calculated via 76GS and KS methods and that between EMT scores using 76GS and MLR methods, whereas a positive correlation should exist between EMT scores from the MLR and KS methods. Indeed, for both NCI-60 and CCLE datasets, the EMT scores calculated via different methods were found to be correlated significantly with a high absolute value of correlation coefficients in the expected direction, when compared pairwise (**Figure 2, S1**). Given that the three scoring methods utilize very different metrics and varying number of genes to define and quantify EMT, it was remarkable that all three showed such high consistency in scoring EMT for these datasets that contained cell lines across different cancer types.

**Figure 2:**
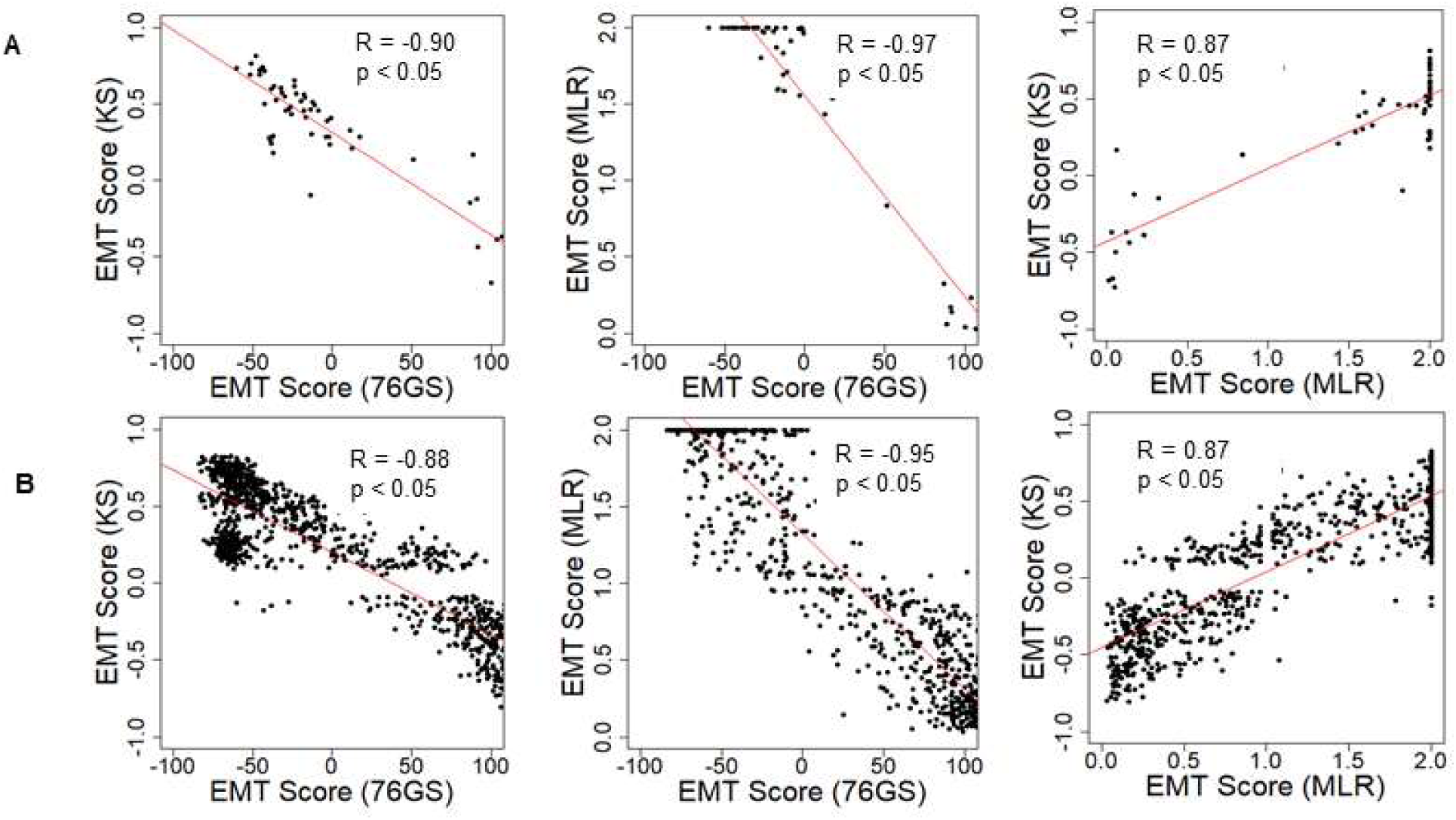
Scatter plot depicting the correlation between the EMT scores of cancer cell line samples, calculated via three methods. Each pairwise relation is estimated by a linear regression line (red), Pearson’s correlation coefficient (R) and p-value (p) reported in each plot. **A)** NCI60 dataset **B)** CCLE dataset

Next, we investigated whether this trend was also present in the TCGA patient samples of different tumor types. Again, the trend remained consistent across tumor types – a strongly positive significant correlation between scores via MLR and KS, and a strongly negative significant correlation between scores via 76GS and KS and those via 76GS and MLR methods (**Figure 3A-C, S3**). Among all tumor types in TCGA data, breast cancer exhibited the highest observed correlation coefficient across methods (**Figure 3C**). Thus, the association between EMT scores and patient survival was assessed using breast cancer patient samples. The samples were scored using all three methods and segregated into EMT^high^ and EMT^low^ groups based on the mean value of each EMT score. The three EMT scoring methods showed consistent trends in predicting overall survival highlighting that patients with a strongly mesenchymal phenotype had better survival probability (**Figure 3D**), endorsing the emerging notion that the predominance of EMT in primary tumors and/or circulating tumor cells (CTCs) need not always be correlated with worse patient survival (Tan et al., 2014; Saxena et al., 2019).

**Figure 3:**
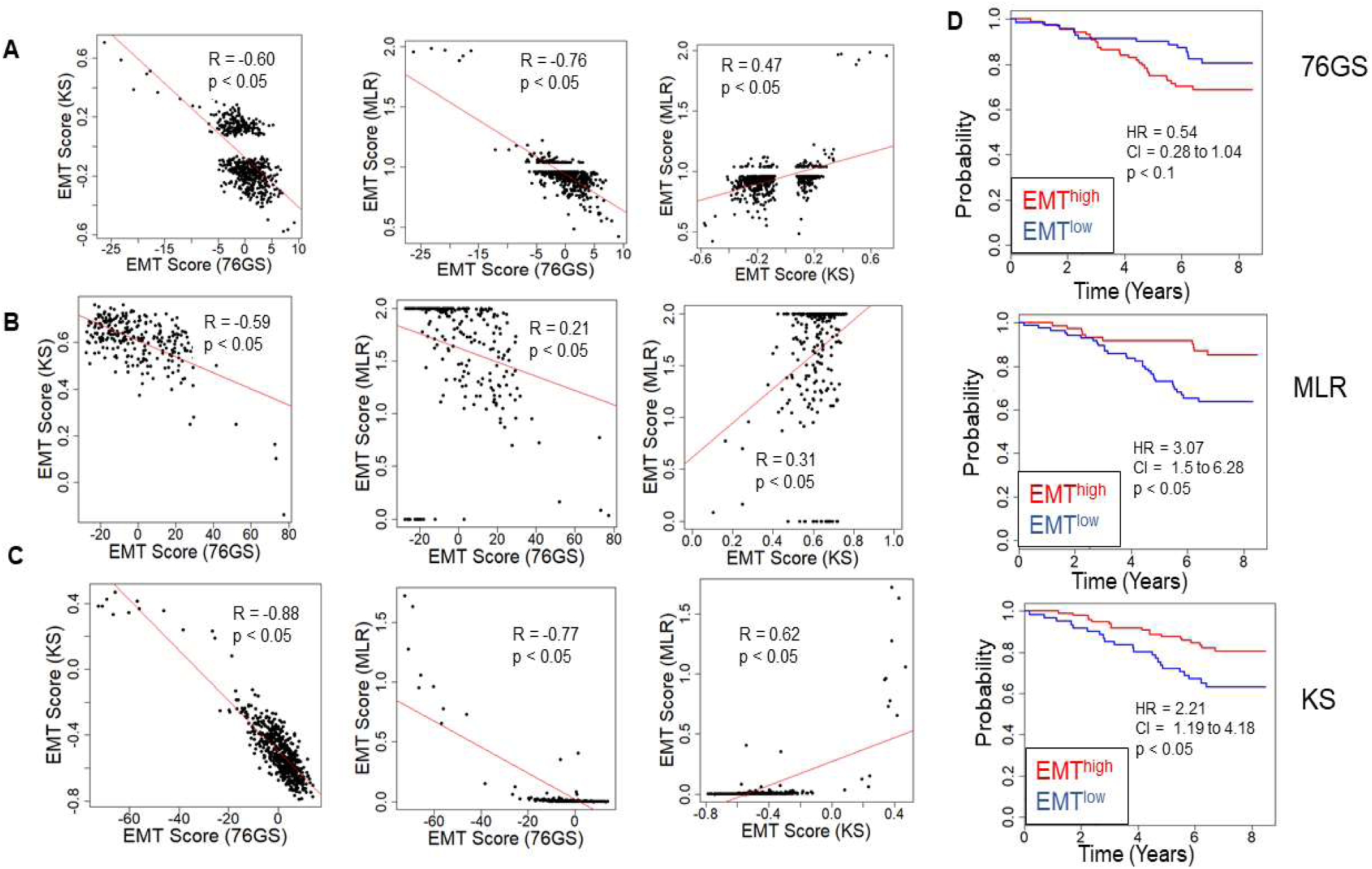
Concordance across all three EMT scoring methods in quantification of EMT and survival prediction of tumor patients. Each pairwise relation is estimated by linear regression (red), Pearson’s correlation coefficient (R) and p-value (p), reported in each plot. **A**) TCGA ovarian cancer dataset, **B**) TCGA sarcoma dataset, **C**) TCGA breast cancer dataset. **D)** Correlation between EMT status (high vs. low) and overall survival (OS) in breast cancer patients. Kaplan–Meier survival analysis is performed to estimate differences in survival of EMT^high^ and EMT^low^ groups of patient samples in GSE1456. P-values (p) reported are based on log rank test. HR (hazard ratio) and CI (95% confidence interval) reported, are estimated using cox regression.

EMT can be driven by diverse biomechanical and/or biochemical stimuli in tumor microenvironments. TGFβ is one of the best-studied drivers of EMT, and a recent study identified a signature specific to TGFβ-induced EMT (Foroutan et al., 2017). EMT scores calculated via any of the three methods – KS, MLR, 76 GS – correlated well with the scores calculated for TGFβ-induced EMT gene signature (**Figure S2**), further endorsing the equivalence of these methods in identifying the onset of EMT.

After establishing this consistency in *in vitro* cell line datasets and TCGA patient samples, we focused on several publicly available microarray datasets including those of EMT induction or reversal, isolation of subpopulations etc. Each dataset comprised a variety of samples in terms of different cell lines, conditions and treatments. An analysis of different GEO datasets showed that EMT scores calculated via these three methods, when compared pairwise, were significantly correlated in the expected direction (**Figure 4A, Table S5**). Out of 85 different datasets, a large percentage of them showed trends in the expected direction (62/85 in KS *vs.* 76GS; 64/85 in MLR *vs.* 76GS; 49/85 in MLR *vs.* KS) (**Figure 4B**). Strikingly, 43 datasets were found to be common across all the pairwise comparisons (**Figure 4C**), establishing a high degree of concordance among EMT scores calculated via these three EMT scoring methods.

**Figure 4:**
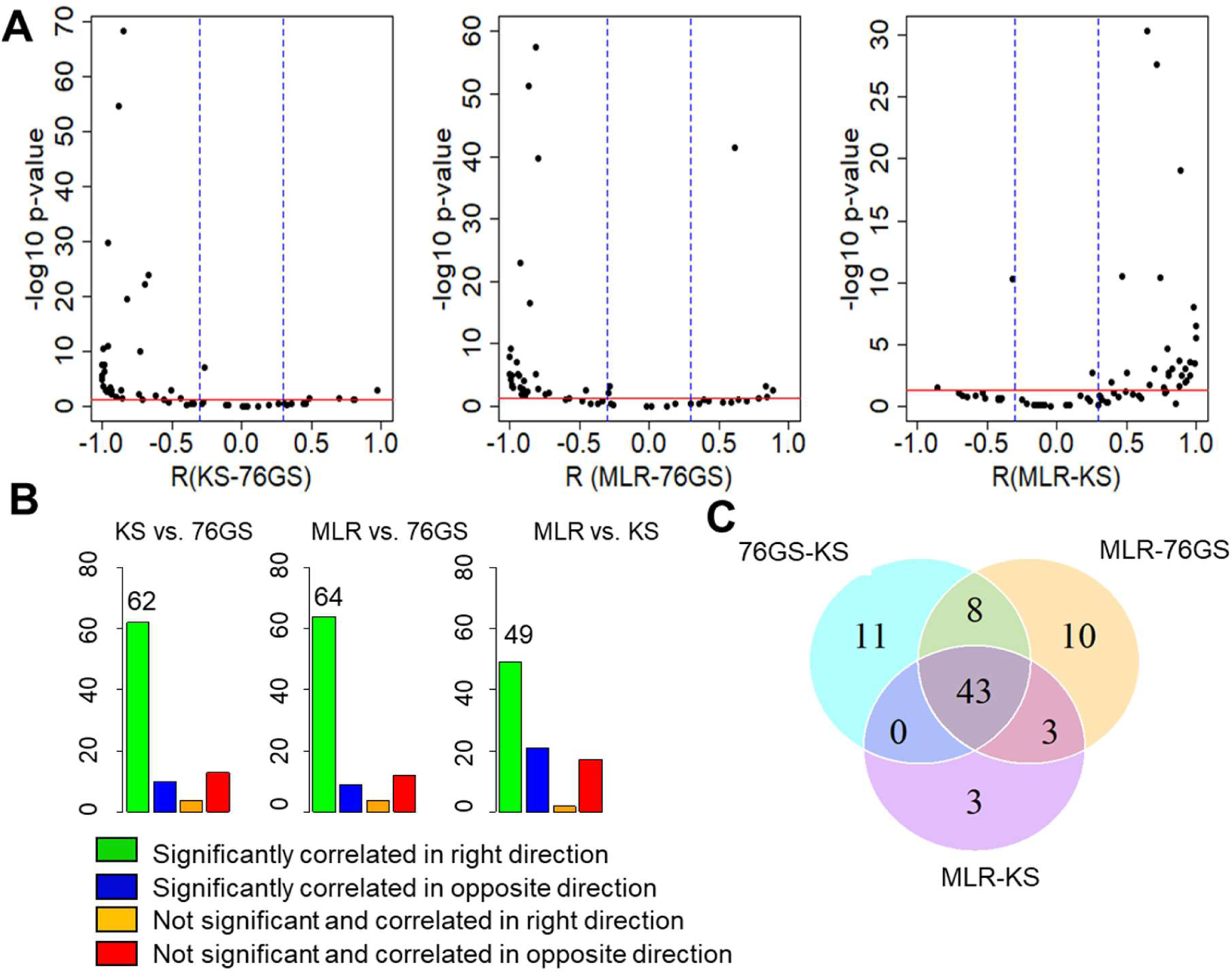
Plots depicting pairwise comparisons of all three EMT scores **A**) Volcano plots showing the correlation of different EMT scoring methods across 85 different GEO microarray datasets along with the p-values for the respective correlation coefficient values. In each case, −log10(p-value) is plotted as a function of Pearson’s correlation coefficient. Thresholds for correlation (R < −0.3 or R > 0.3; vertical blue lines) and p-values (p <0.05; horizontal red line) are denoted. **B)** Bar plots for different categories based on the correlation sign and statistical significance of all three pairwise comparisons across 85 datasets**. C)** Venn diagram showing the common GEO datasets across all pairwise comparisons that are significantly correlated in the expected direction.

Next, we investigated specific cases where EMT/MET was induced in various cell lines by different EMT/MET regulators. Lung cancer cell lines A549, HCC827 and H358 in which EMT was induced by TGFβ showed higher EMT scores using MLR and KS methods, but lower scores via 76GS method, as compared to untreated ones (**Figure 5A**). Similarly, the epithelial breast cancer cell line MCF-7 transfected to overexpress EMT-inducing transcription factor Snail exhibited a more mesenchymal phenotype relative to the control, as identified via all three scoring methods (**Figure 5B**). Consistent trends were seen in EpRAS tumor cells treated with TGFβ (**Figure 5C**), and in human mammary epithelial cells HMLE overexpressing one of the three EMT-inducing transcription factors (EMT-TFs) – SNAI1 (Snail), SNAI2 (Slug), and TWIST (**Figure 5D**). Interestingly, all three scoring methods suggested that EMT induced by Snail or Slug was stronger than that induced by Twist (**Figure 5D**). Further, inducing EMT via overexpression of EMT-TFs Twist, Snail, Goosecoid, or treatment with TGFβ or knockdown of E-cadherin was capable of altering the EMT scores of HMLE cells (**Figure S4C**).

**Figure 5:**
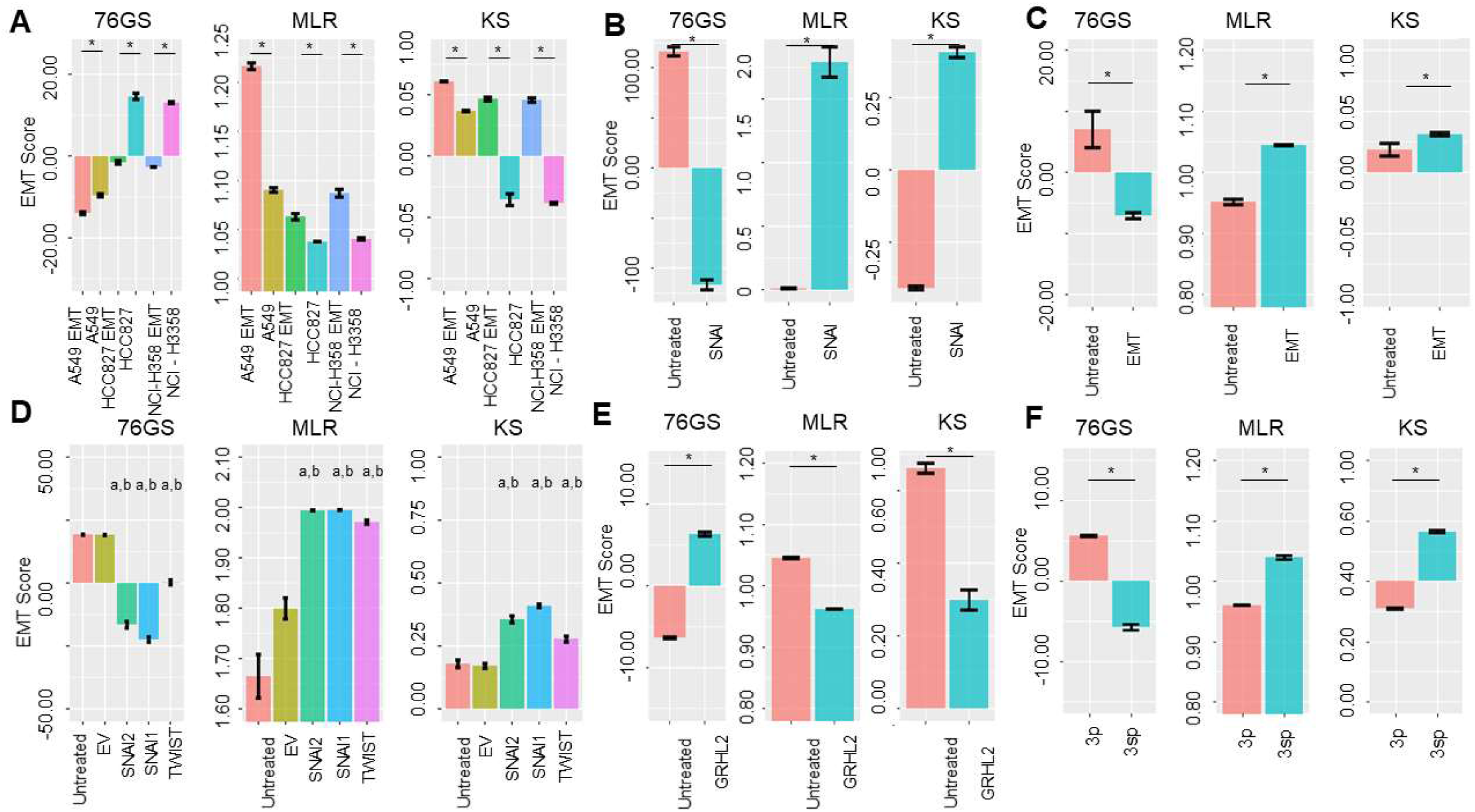
Bar plots showing EMT scores of different cell lines calculated using the three EMT scoring methods. **A)** EMT induction is shown in three cell lines – A549, HCC87 and NCIH358 (GSE49664). **B)** EMT induction in MCF7 cell line (GSE58252). **C)** EMT induction in EpRas cells (GSE59922). **D)** EMT induction by different EMT-inducing transcription factors. ‘a’ denotes statistical significant difference (p<0.05, n=3, two-tailed Students’ t-test) for pairwise comparison of a given set with untreated (first column), ‘b’ denotes the same when compared with empty vector (EV; second column). (GSE43495). **E)** MET induction by GRHL2 in MDA-MB-231 cell line (GSE36081). **F)** Two cell lines of hepatocellular carcinoma with varying EMT status (GSE26391). Each control case has been compared to EMT/MET induced case (* p< 0.05, n=3, two-tailed Student’s t-test; error bars represent standard deviation)

Additionally, these three methods also captured the reversal of EMT – Mesenchymal-Epithelial Transition (MET) – induced by MET-inducing transcription factor GRHL2 in MDA-MB-231 cells (**Figure 5E**). Moreover, baseline differences in EMT status between two hepatocellular carcinoma cell lines identified experimentally (Van Zijl et al., 2011) were also recapitulated by all three scoring methods; while HCC-1.2 (referred to as 3p) showed more epithelial features, HCC1.1 (referred to as 3sp) was relatively more mesenchymal (**Figure 5F**). We also calculated the EMT scores for the dynamic EMT time series datasets (i.e. cases where more than 2 time points were available for EMT induction); all three methods were able to recapitulate the relevant trends in EMT scores as expected when EMT was induced in A549 or LNCAP cells (**Figure S4A, D**). Finally, we calculated EMT scores for a population of circulating tumor cells (CTCs) collected from breast cancer patients, and observed heterogeneity in CTCs along the epithelial-hybrid-mesenchymal spectrum (**Figure S4B**), thus reminiscent of similar observations based on immunohistochemical staining of a few canonical markers (Yu et al., 2013).

### Variability in EMT scores measures tumor heterogeneity

Recent studies have emphasized that intra-tumor heterogeneity in patients and inter-tumor heterogeneity in a given cancer subtype can accelerate progression and metastasis (Lawson et al., 2018). Thus, we were interested in identifying which tumor types are more heterogeneous with regard to EMT scores calculated via the three methods. We grouped the CCLE samples by different tumor types and calculated the mean and variance of all EMT scores across a given tumor. The EMT scores, calculated across the three methods, showed less variation in the EMT scores of the tumor types of mesenchymal origin such as sarcoma and lymphoma, as compared to that of the other tumor types such as breast cancer and lung cancer (**Figure 6A-C, Table S7**). The most heterogeneous tumor types identified based on the variance in EMT scores largely overlapped for all methods: a) breast cancer, b) stomach cancer, c) non-small cell lung cancer, d) bile duct cancer, and e) urinary tract cancer (**Figure 6A-C**). We also calculated pairwise correlations of EMT scores across all the tumor types and observed consistently significant trends (**Table S8**).

**Figure 6:**
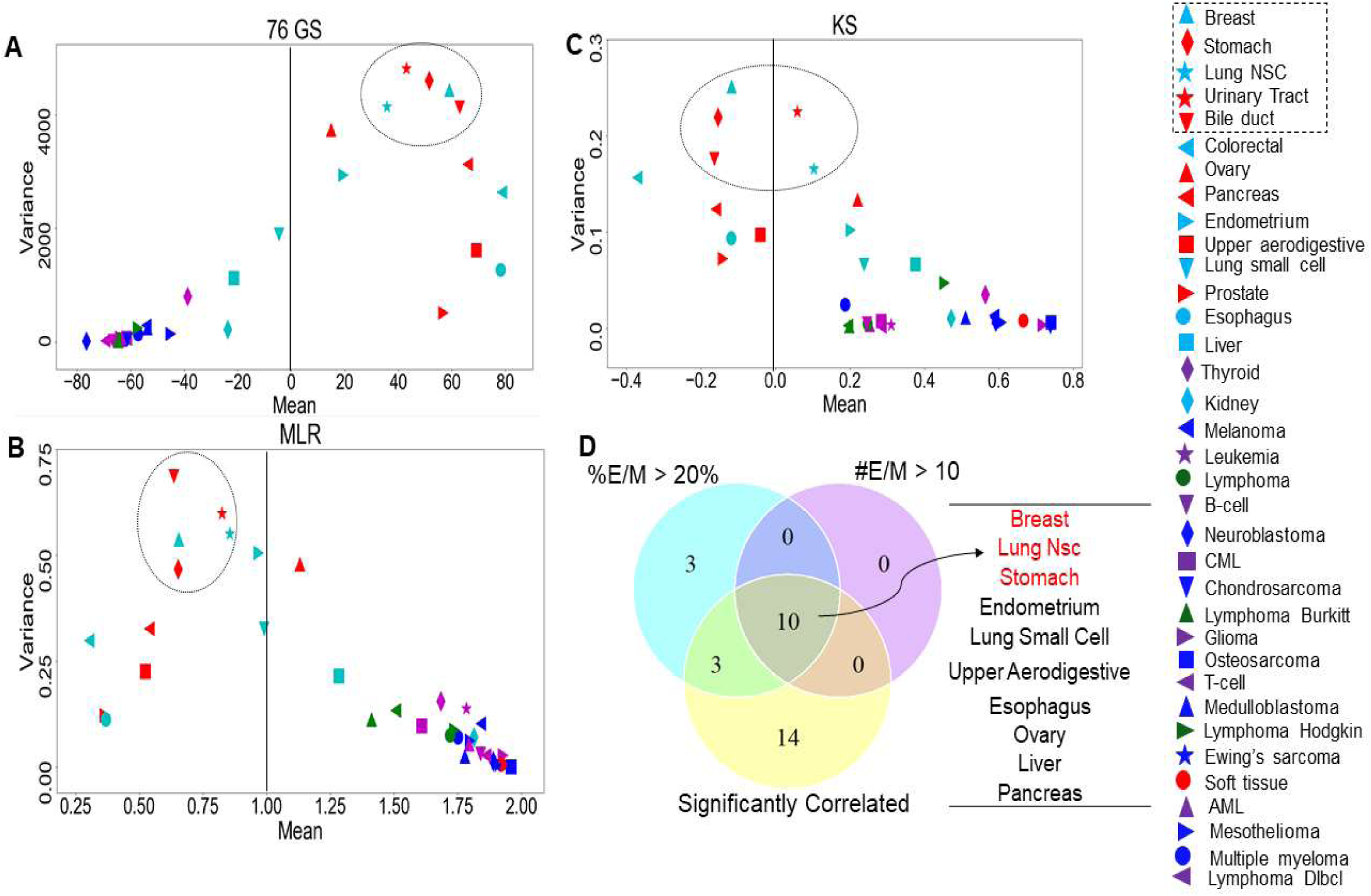
Variance and mean of EMT scores in CCLE samples grouped by tumor subtype, highlighting the most variable tumor types (circled). **A)** 76GS EMT Scores **B)** MLR EMT Scores **C)** KS EMT Scores. **D)** Venn diagram showing the overlap between the tumor types based on the abundance of hybrid samples as defined by the MLR method, where #EM > 10 denote the cases where the absolute number of hybrid E/M samples in a tumor subtype is more than 10; %EM > 20 denote the cases where the percentage of cell lines identified as hybrid E/M in a given tumor subtype is greater than 20%.

One of the proposed mechanisms underlying such heterogeneity in EMT status has been epithelial-mesenchymal plasticity, i.e. the proclivity of individual cells in a population to obtain and switch among multiple phenotypic states. Such plasticity is typically seen to be higher in cells in one or more hybrid E/M states (Pastushenko and Blanpain, 2019; Tripathi et al., 2019b, 2019a). Thus, we asked whether the frequency of hybrid E/M phenotype contributes to heterogeneity in terms of EMT scoring. One of the EMT scoring methods – MLR – calculates the probability of a given transcriptomic profile being associated with the epithelial, hybrid E/M or mesenchymal state, thus enabling us to identify hybrid E/M samples specifically. First, we found that the variance of EMT scores was the highest in samples identified as hybrid E/M **(Table S6)**. Next, we checked the relative frequency and absolute number of hybrid E/M samples (as defined by MLR method) across tumor types, among the cases where EMT scores calculated via all three methods were significantly correlated. Indeed, the tumor types that met the three conditions – a) total number of hybrid E/M samples being more than 10, b) percentage of hybrid E/M samples being greater than 20%, and c) a good correlation among all three methods – were enriched in the most variable tumor types (**Figure 6D**), suggesting hybrid E/M phenotypes contribute maximally to E-M heterogeneity (**Table S9**).

We also calculated the correlations in EMT scores obtained from each method, after segregating the cell line samples into E, E/M and M, based on predictions from the MLR method. The correlation coefficients within the E, E/M and M subgroups of a given tumor subtype were observed to be somewhat different than those found for all tumor subtype samples without any partitioning into E, E/M and M subgroups (**Table S8**). These results suggest that while a generic trend in terms of EMT scores is seen across the three methods, the categorization in terms of E, E/M and M may vary to some degree based on the EMT scoring method used.

### Individual hybrid E/M samples are different from hybrid mixtures of E and M

A given transcriptomic profile may be classified as hybrid E/M for several reasons: a) the sample contains individually hybrid E/M cells (hybrids), b) the sample contains a mixture of epithelial and mesenchymal cells (mixtures), or c) the sample contains a combination of hybrids and mixtures. We sought to distinguish true hybrids from mixtures based on an additional feature of MLR scoring - mixture curve analysis (Jia et al., 2019). This analysis quantifies the distance of a given sample from a ‘mixture curve’ which connects the position of mean signatures of ‘pure’ epithelial and ‘pure’ mesenchymal samples. The farther a given sample is from the mixture curve, the higher the likelihood of that particular sample containing truly hybrid E/M cells.

First, we determined the mixture curves based on the CCLE samples. We ranked all cell lines in the CCLE dataset based on their EMT scores and identified the top 35 most epithelial (i.e. lowest 35 in terms of MLR EMT scores) and top 35 most mesenchymal samples (i.e. highest 35 in terms of EMT MLR scores). Then, the mixture curve was determined based on the convex combinations of mean signatures of these 35 ‘pure’ E and 35 ‘pure’ M reference samples. All the CCLE cell lines identified as hybrid E/M were then plotted alongside the mixture curve (**Figure 7A**) and their distances from the curve were calculated. While some samples fell close to the curve, many deviated substantially (**Figure 7B**). We subsequently picked the farthest and the closest 10, 20, 50 and 100 samples from the mixture curve and calculated their EMT scores. Intriguingly, the mean EMT score of samples farthest from the mixture curve was different than that of the closest samples as calculated using MLR (**Figure 7C**). This trend persisted despite varying the number of samples and EMT scoring method utilized for analysis (**Figure S5A,B**). Similarly, another ‘mixture curve’ based on median of 35 ‘pure’ E and ‘pure’ M reference samples was obtained from CCLE dataset; all trends observed previously for mixture curve based on the mean were recapitulated (**Figure S5C-F**). In each case (mean or median mixture curve; across the three EMT scoring methods; N=10, 20, 50, 100), the cell lines closest to the mixture curve tended to be more mesenchymal than the ones farthest from the curve (**Figure 7C, S5**).

**Figure 7:**
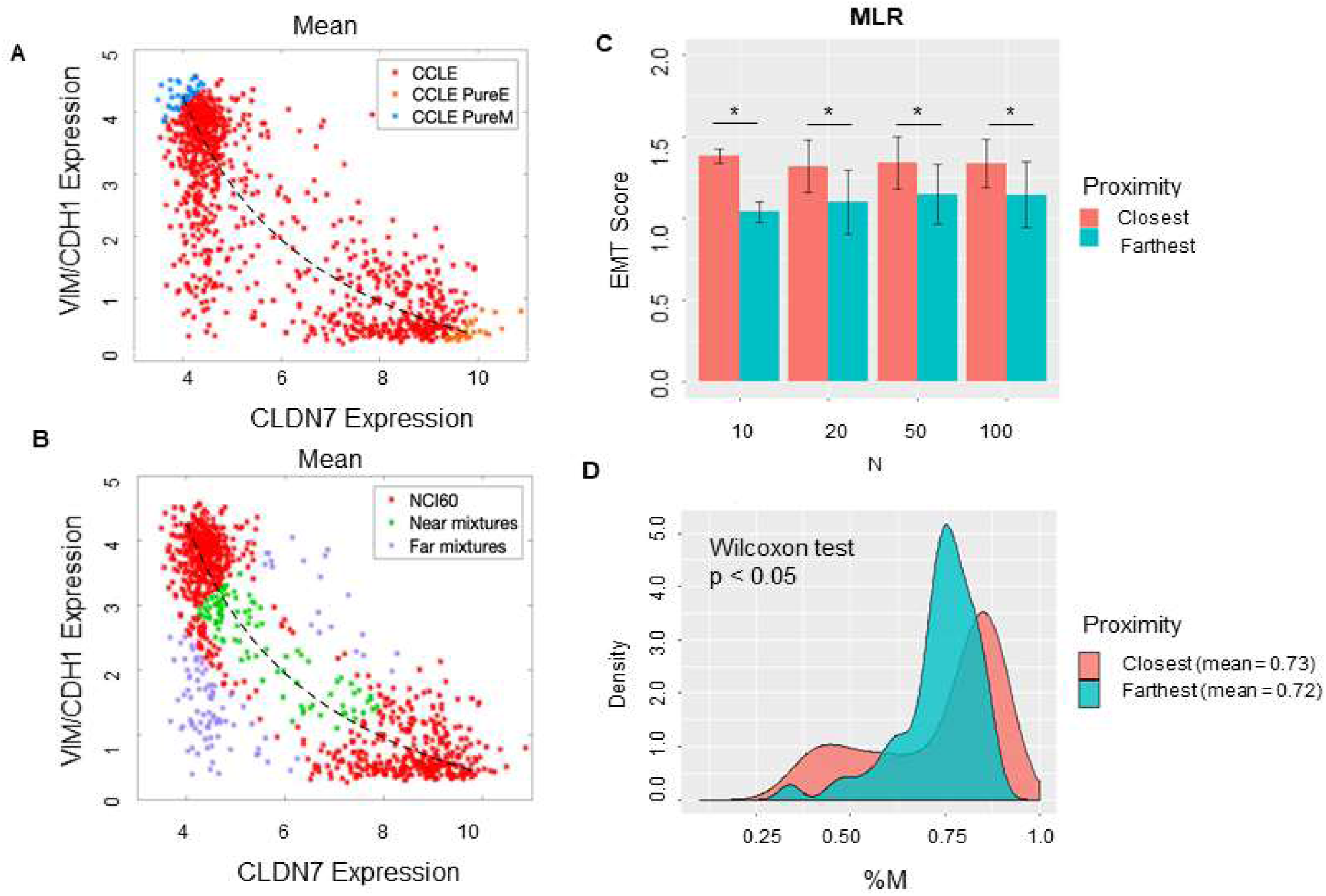
Distinguishing between hybrid E/M cells vs. mixtures of E and M cells. **A)** Scatter plot showing CCLE cell lines that display a hybrid E/M phenotype (red) on the mixture curve (dotted curve) determined by the mean of 35 pure E (orange) and pure M (blue) reference samples in CCLE dataset. **B)** Scatter plot showing the 100 farthest (purple) and 100 closest (green) samples based on the distance from the mixture curve. **C)** Bar plots showing EMT scores of N (10, 20, 50, 100) closest and farthest hybrid E/M samples from mixture curve. **D)** Mesenchymal proportion (%M) distribution of the 100 closest and farthest hybrid samples from mixture curve.

In order to distinguish the hybrid E/M samples from mixtures of pure E and pure M samples, we lastly characterized the composition of the closest and farthest hybrid E/M samples by estimating the percentage of mesenchymal phenotype (%M) in each sample based on the convex combination ‘mixture curve’ in the two-dimensional space (VIM/CDH1 expression; CLDN7 expression). While the difference in mean values of the composition (%M) of closest and farthest samples, was marginal, but their overall distributions in terms of %M differed substantially. The closest samples exhibited a bimodal distribution, while the farthest samples showed a broad-tailed distribution (**Figure 7D**). This analysis demonstrates the possibility of a quantifiable compositional difference between truly hybrid E/M samples and mixtures of E and M cells.

## Discussion

EMT is a reversible and dynamic process which has been shown to be activated during cancer progression. EMT involves a multitude of changes at both molecular and morphological levels. Various attempts to characterize the spectrum of EMT at molecular and/or morphological levels have been made recently, enabled by latest developments in multiplex imaging, single-cell RNA-seq and inducible systems (Mandal et al., 2016; Pastushenko et al., 2018; Stylianou et al., 2018; Cook and Vanderhyden, 2019; Devaraj and Bose, 2019; Karacosta et al., 2019; Wang et al., 2019b; Watanabe et al., 2019). These approaches have highlighted the dynamical nature of EMT in driving cancer progression in patients (Jolly and Celia-Terrassa, 2019). Further, various approaches to quantify the EMT spectrum of samples based on different signatures of tumor types have been made (Foroutan et al., 2017; Puram et al., 2017). Among all the methods available for EMT scoring, we have compared the ones that are more generalized – KS (Tan et al., 2014), MLR (George et al., 2017), and 76 GS (Byers et al., 2013; Guo et al., 2019). These three methods use different combinations of genes and metrics; however, they show a very good concordance amongst them in terms of identifying an empirical trend along the EMT axis.

Here, we compared the aforementioned EMT scoring metrics for their ability to identify the onset and extent of EMT/MET via calculating EMT scores for cell line cohorts NCI-60 and CCLE, TCGA cohorts from multiple subtypes, and datasets containing samples with overexpression and/or knockdown of many EMT/MET inducers such as TGFβ, Snail, Slug, Twist, E-cadherin, GRHL2 (De Craene and Berx, 2013). The remarkable concordance among the EMT scores calculated via the methods analysed above suggests some macroscopic signal that can resolve the extent of EMT in a given sample amidst the complexity of EMT and the networks regulating it. It is plausible that within these regulatory networks, there exist key nodes forming one (or more) core circuit(s) which receive(s) a large number of inputs and may have diverse outputs, reminiscent of bow-tie structures seen in biological networks of cell-fate decision-making (Friedlander et al., 2015). This idea of core circuit(s) driving EMT is substantiated by transcriptomic meta-analysis identifying common signatures for EMT driven by distinct inducers (Taube et al., 2010; Liang et al., 2016). For instance, one network motif commonly found in core circuits regulating EMT and associated traits is a mutually inhibitory feedback loop between two ‘master regulators’ driving opposing cell phenotypes (Hong et al., 2015; Huang et al., 2015; Saha et al., 2018); for instance, ZEB1 driving EMT and miR-200 driving MET (Jia et al., 2017). An intricate coupling among such feedback loops may give rise to a spectrum of EMT phenotypes as has been seen across cancer types in cell lines, circulating tumor cells and primary tumor biopsies (Armstrong et al., 2011; Huang et al., 2013; Schliekelman et al., 2015; Andriani et al., 2016; Iyer et al., 2019; Markiewicz et al., 2019; Varankar et al., 2019).

As a consequence of computing the extent of EMT in a given transcriptomic sample, these three methods were able to identify the most variable tumors in terms of their EMT status. Most tumors of mesenchymal lineage such as sarcomas were shown to be least variable, i.e. CCLE cell line belonging to these tumors were identified as quite mesenchymal and similar among themselves, whereas breast cancer, non-small cell lung cancer, bile duct cancer, urinary tract cancer, and stomach cancer exhibited the largest degree of variability in terms of their inherent EMT status. The observations about sarcomas, breast cancer, and non-small cell lung cancer are well-supported by existing experimental data (Blick et al., 2008; Schliekelman et al., 2015; Jolly et al., 2019b); however, the predictions about other cancer types identified to be more heterogeneous need further investigation. Our results also demonstrate a link between the predominance of hybrid E/M status and heterogeneity patterns, possibly emerging due to relatively higher plasticity of cells in one or more hybrid E/M phenotypes (Pastushenko et al., 2018; Tripathi et al., 2019b). These observations also imply that tumor types with a greater number of hybrid E/M cells may require alternative treatment strategies as compared to those containing predominantly epithelial or predominantly mesenchymal populations.

This comparative analysis of the three methods shows two key advantages of MLR method. First, it uses the least number of genes to calculate an EMT score - 23 genes required by MLR as compared to 76 genes by 76GS, and 315 genes for tumor and 218 genes for cell lines by KS. This feature is important because 23 genes can be relatively easily measured experimentally without microarray or RNA-seq. Second, the MLR method, by virtue of its underlying theoretical framework, is capable of isolating hybrid E/M samples and has been expanded to identify whether the resultant gene expression is more likely to derive from ‘true’ individual hybrid E/M samples or admixtures of epithelial and mesenchymal samples. While, in theory, other methods could adopt similar adaptations to address this issue in the future, the resolution of E, M, and hybrid E/M populations through those methods would require analysing a higher dimensional subspace of the original predictors, given the large number of genes used by those methods to calculate EMT scores as compared to MLR method. This contrasts with the MLR method, where the mixture analysis is performed directly on the two-dimensional EMT predictor space (CLDN7, VIM/CDH1) utilized by this method. Distinguishing between these possibilities is critical because the behavior of mixtures of epithelial and mesenchymal samples vs. truly hybrid E/M samples can be strikingly different; a recent study showed that the presence of hybrid E/M cells is essential to form tumors in mice, a task which could not be achieved as efficiently by co-cultures of epithelial and mesenchymal cells alone (Kröger et al., 2019).

Our analysis shown here suffers from following limitations. First, in terms of classifying hybrid E/M into ‘pure’ hybrid E/M vs. mixtures of epithelial and mesenchymal subpopulations, we have considered mutually exclusive criteria: a) a sample identified as hybrid E/M at a bulk level contains mixtures of epithelial and mesenchymal subpopulations, and b) a sample identified as hybrid E/M at a bulk level contains all ‘true’ hybrid E/M cells. However, many cell lines may contain cells in each of the three phenotypes in varying ratios (Ruscetti et al., 2016; George et al., 2017; Jia et al., 2019). Thus, future efforts should aim to identify the relative proportions of these three different phenotypes in a given sample. Second, although we show that among the samples identified to be lying closest vs. farthest from the ‘mixture curve’ by MLR, all three EMT scoring metrics suggested that the ones lying closest to the curve are more mesenchymal than the ones lying farthest from the same, we lack a clear biological interpretation of this observation. Future efforts will focus on comparing the morphological and functional behavior of the CCLE cell lines identified to be closest vs. farthest from the ‘mixture curve’ generated based on the CCLE samples.

## Acknowledgement

This work was supported by Ramanujan Fellowship (SB/S2/RJN-049/2018) awarded by Science and Engineering Research Board (SERB), Department of Science and Technology (DST), Government of India. HL was supported by National Science Foundation (NSF) grants PHY-1427654 (Center for Theoretical Biological Physics) and PHY-1935762. JTG was supported by National Cancer Institute of NIH (F30CA213878).

## Supplementary Figures

**Figure S1:**
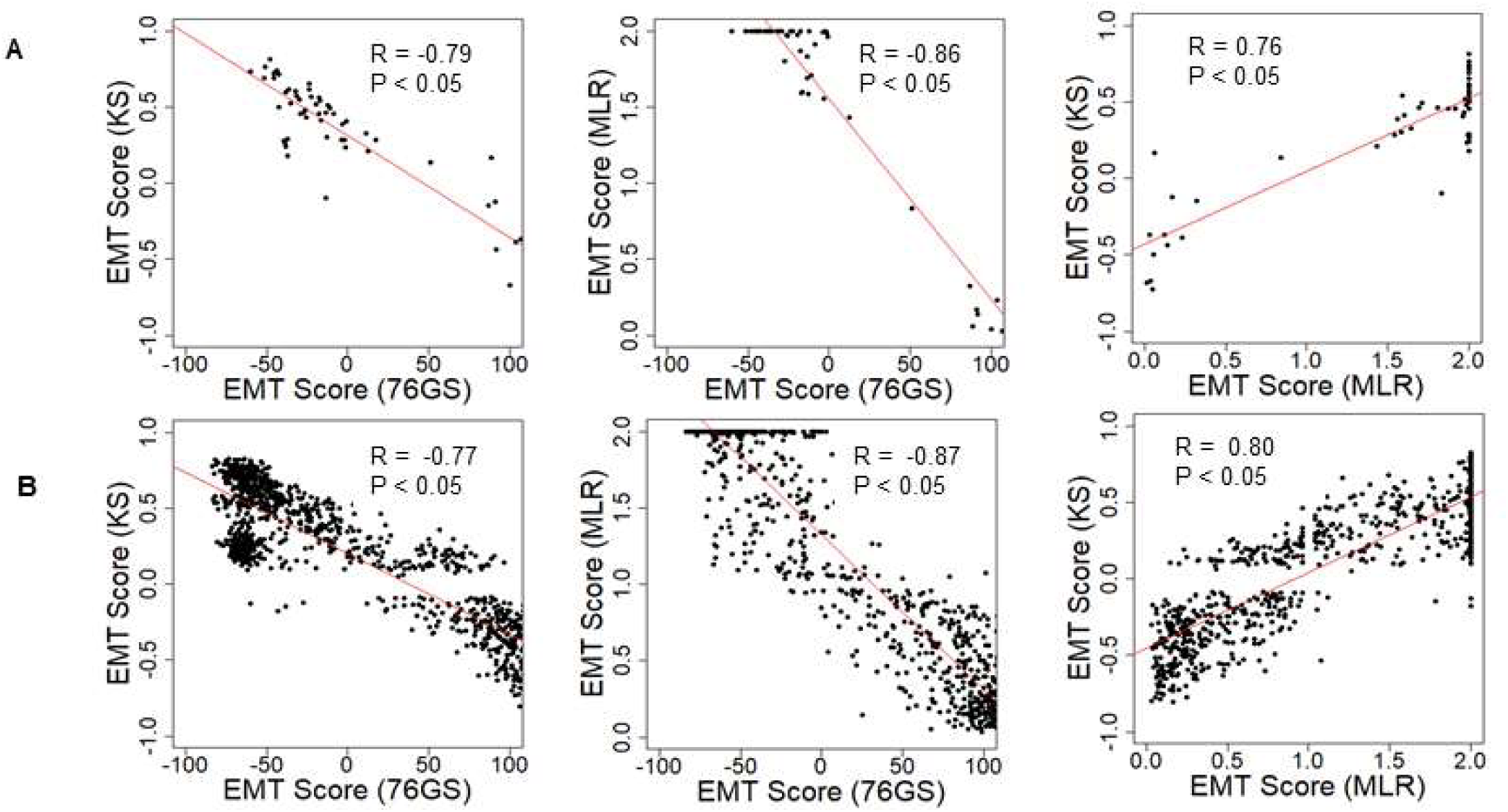
Scatter plot depicting the correlation between the EMT scores of cancer cell line samples, calculated via three EMT scoring methods. Each pairwise relation is estimated by a linear regression line (red), Spearman’s correlation coefficient (R) and p-value (p) reported in each plot. **A)** NCI60 dataset **B)** CCLE dataset

**Figure S2:**
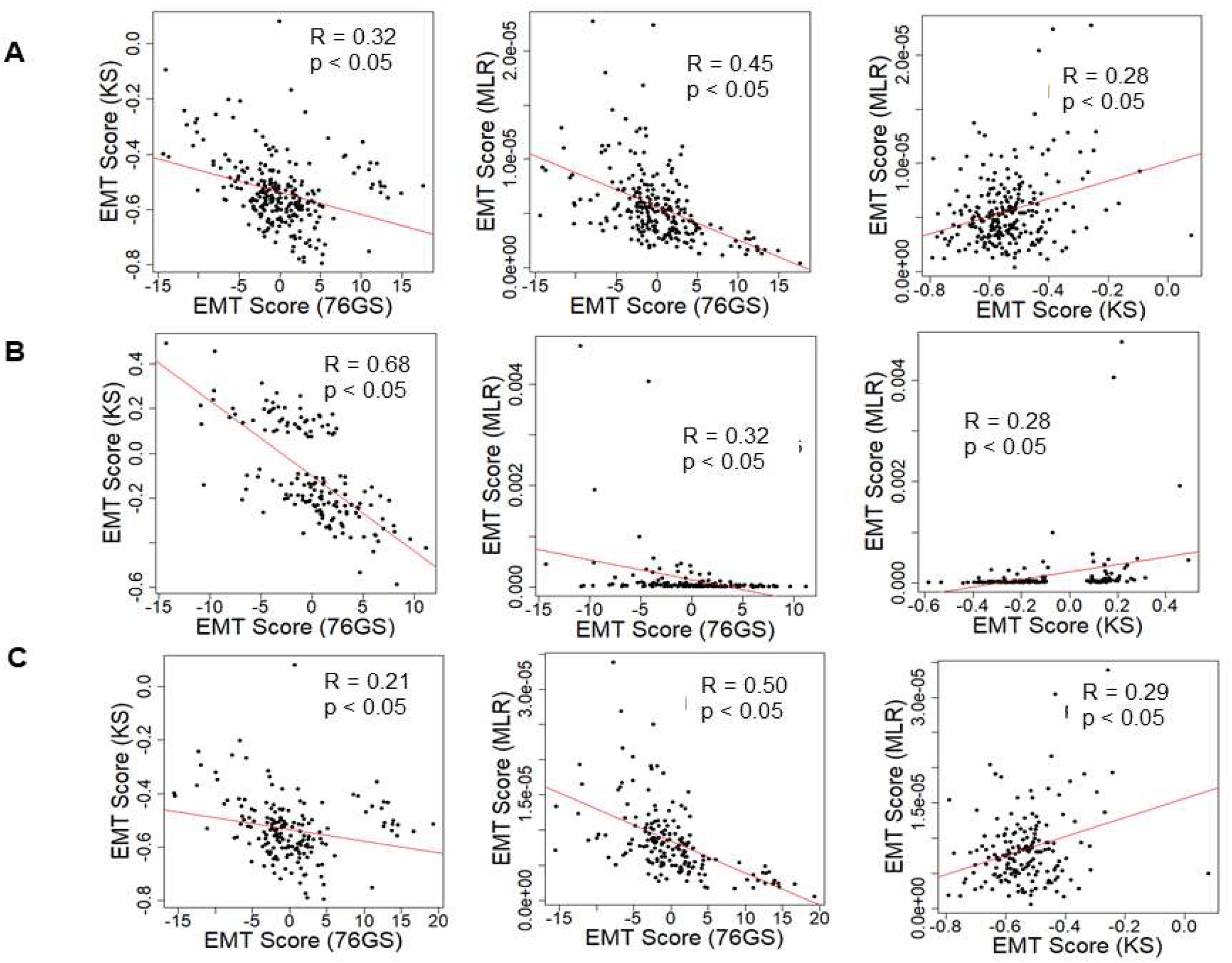
Scatter plot depicting the correlation between the EMT scores of different tumor types in TCGA dataset, calculated via three methods. Each pairwise relation is estimated by a linear regression line (red), Pearson’s correlation coefficient (R) and p-value (p) reported in each plot. (A) Lung squamous cell cancer (B) Colon adenocarcinoma (C) Colon and rectal adenocarcinoma

**Figure S3:**
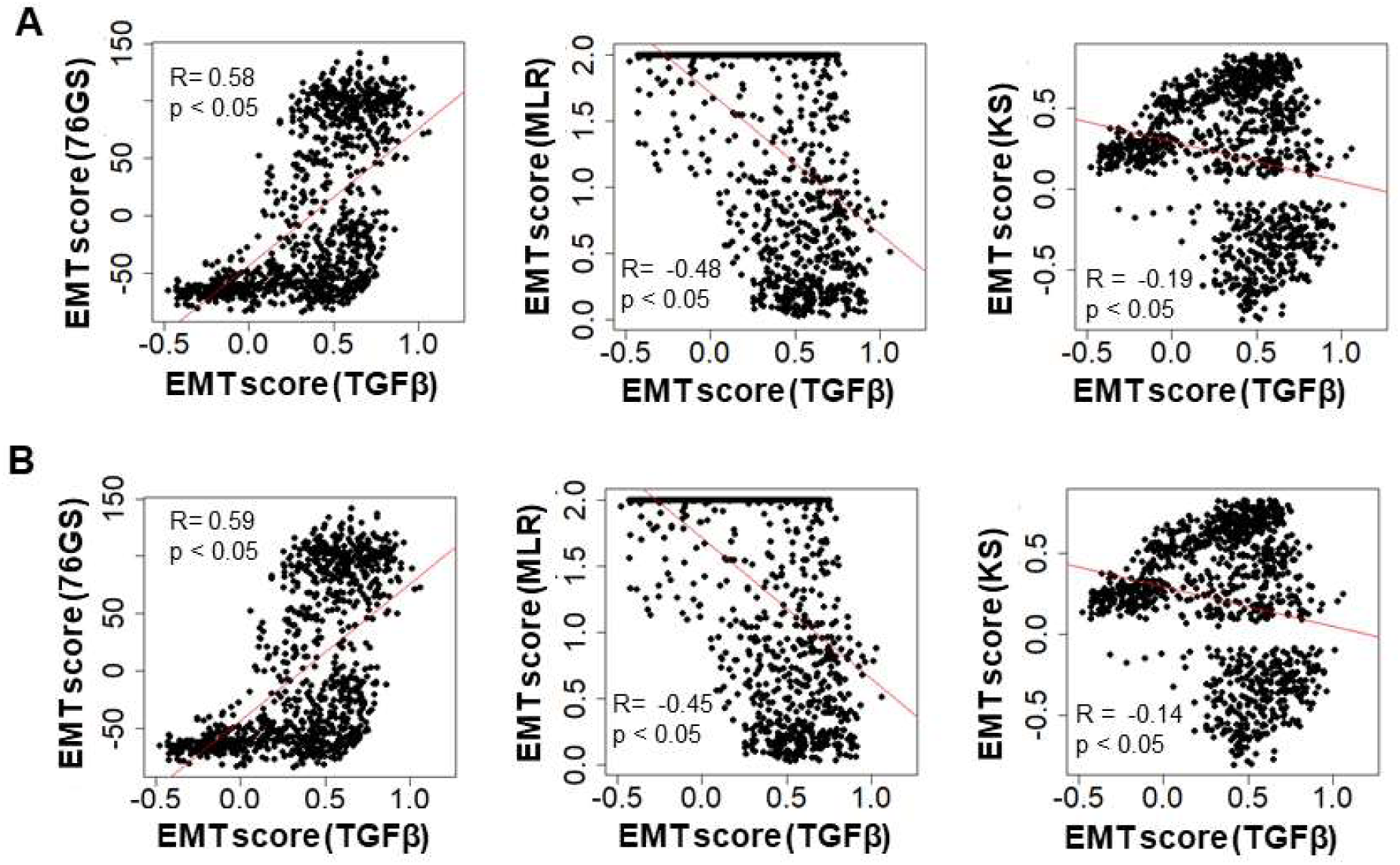
EMT score correlation with TGFβ specific EMT scoring method in CCLE dataset(A) Pearson’s correlation coefficient (B)Spearman’s correlation. Correlation coefficient (R) and p-value (p) reported in each plot

**Figure S4:**
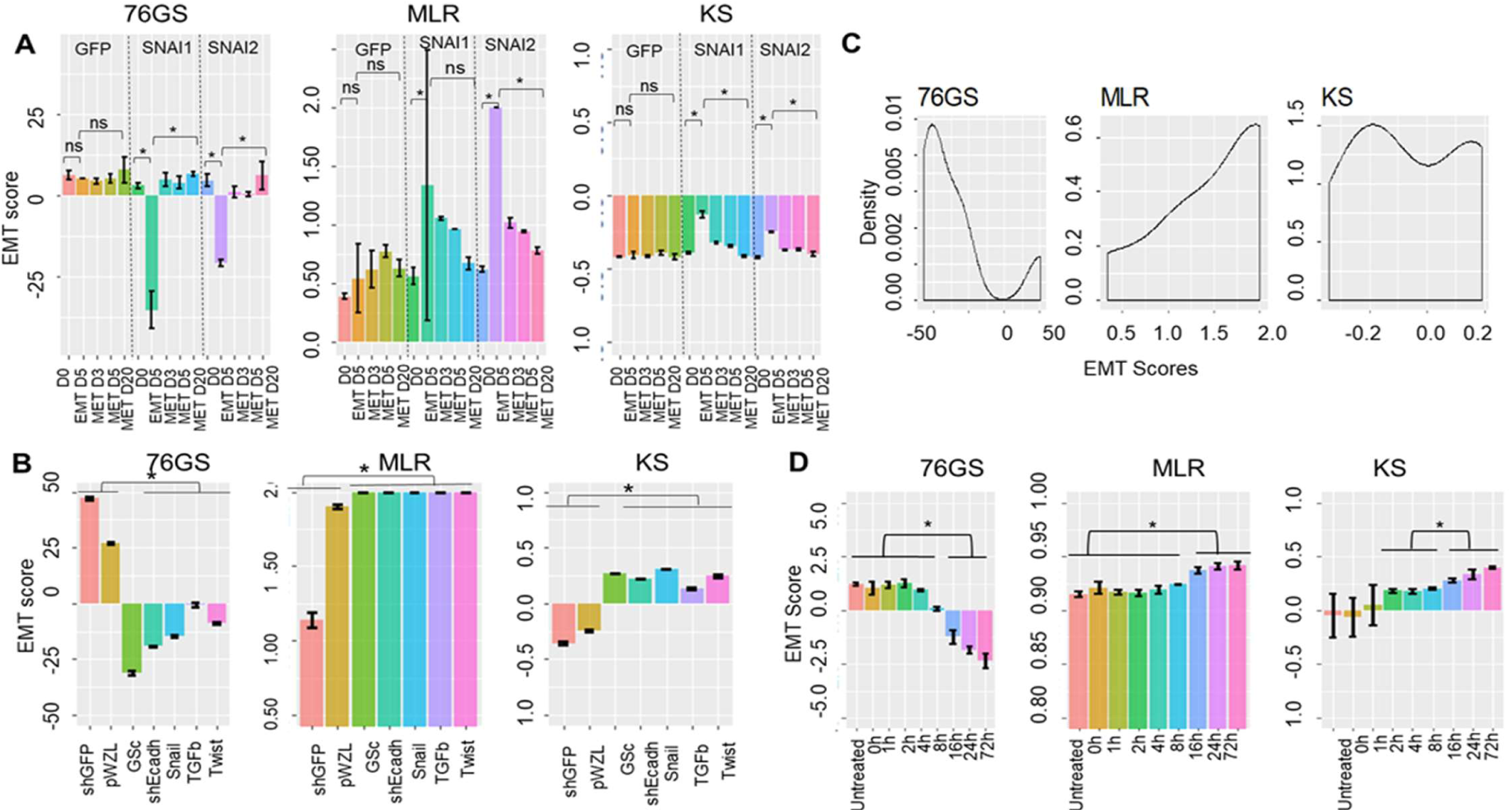
Bar plots showing EMT scores of different EMT time series datasets (A) GSE84002-EMT and MET induction over time by GFP, SNAI1 and SNAI2. (B) GSE24202-EMT induction by different EMT regulators (C) Tumor CTCs (D) GSE17708 – EMT induction over time. (* p < 0.05, n=3, two-tailed Student’s t-test; error bars represent standard deviation)

**Figure S5:**
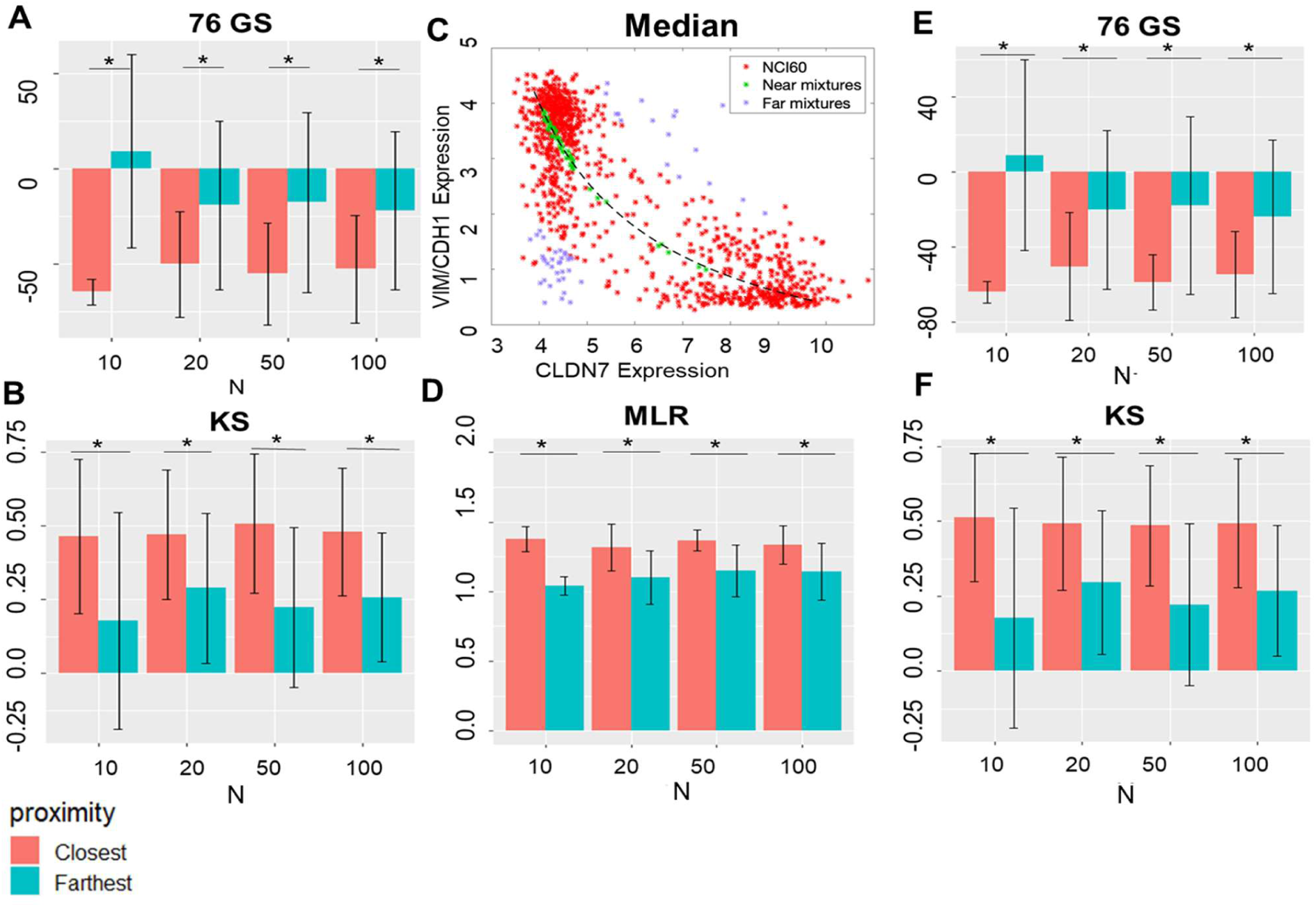
Bar plots showing EMT scores of N (10, 20, 50, 100) closest and farthest hybrid samples from mixture curve. (A) 76GS EMT score (B) KS EMT score (C) Scatter plot showing 100 farthest and closest samples based on the distance from mixture curve defined by median of 35 most pure E and pure M CCLE samples. Bar plots showing EMT scores of N (10, 20, 50, 100) closest and farthest hybrid samples from median mixture curve. (D) MLR EMT score (E)76GS EMT score (F) KS EMT score. (* p < 0.05, N=10, 20, 50 & 100, two-tailed Student’s t-test; error bars represent standard deviation)

## Supplementary Tables

**Table S1:** 76 gene signatures

**Table S2:** List of predictors and normalizers used for calculation of EMT using MLR method

**Table S3:** Epithelial and mesenchymal signature used in KS-statistic (tumor signature)

**Table S4:** Epithelial and mesenchymal signature used in KS-statistic (cell line signature)

**Table S5**: EMT score correlation in list of 85 microarray GEO datasets.

**Table S6**: Mean and standard deviation of EMT score in CCLE samples categorized into E, E/M and M as defined by MLR EMT scores

**Table S7:** Most variable and least variable tumor types based on the coefficient of variation of EMT scores

**Table S8:** Pairwise correlation between all three EMT scores in subcategories (E, E/M and M) across all tumor types of CCLE data

**Table S9:** Abundance of hybrid in different tumor types

